# SQuIRE: Software for Quantifying Interspersed Repeat Elements

**DOI:** 10.1101/313999

**Authors:** Wan R. Yang, Daniel Ardeljan, Clarissa N. Pacyna, Lindsay M. Payer, Kathleen H. Burns

## Abstract

Transposable elements are interspersed repeat sequences that make up much of the human genome. Conventional approaches to RNA-seq analysis often exclude these sequences, fail to optimally adjudicate read alignments, or align reads to interspersed repeat consensus sequences without considering these transcripts in their genomic contexts. As a result, repetitive sequence contributions to transcriptomes are not well understood. Here, we present Software for Quantifying Interspersed Repeat Expression (SQuIRE), an RNA-seq analysis pipeline that integrates repeat and genome annotation (RepeatMasker), read alignment (STAR), gene expression (StringTie) and differential expression (DESeq2). SQuIRE uniquely provides a locus-specific picture of interspersed repeat-encoded RNA expression. SQuIRE can be downloaded at (github.com/wyang17/SQuIRE).

## Introduction

Transposable elements (TEs) are self-propagating mobile genetic elements. Their insertions have resulted in a complex distribution of interspersed repeats comprising almost half of the human genome (Lander et al. 2001; Kazazian 2004). They propagated through either DNA (‘transposons’) or RNA intermediates (‘retrotransposons’)(Huang et al. 2012; Burns and Boeke 2012). Retrotransposons are further classified into Orders based on the presence of long terminal repeats (LTR retrotransposons) or whether they were long or short interspersed elements (LINEs and SINEs)(Wicker et al. 2007). Although most TEs have lost the capacity for generating new insertions over their evolutionary history and are now fixed in the human population, a subset of younger subfamilies from the LINE-1 superfamily (i.e., L1PA1 or L1HS) (Beck et al. 2011), the SINE *Alu* superfamily (e.g., *Alu*Ya5, *Alu*Ya8, *Alu*Yb8, *Alu*Yb9) (Deininger 2011), and composite SVA (SINE-variable number tandem repeat (VNTR)-*Alu*) elements (Hancks et al. 2010) remain retrotranspositionally active and generate new polymorphic insertions (Stewart et al. 2011; Abecasis et al. 2012).

Due to the repetitive nature of TEs, short-read RNA sequences that originate from one locus can ambiguously align to multiple copies of the same subfamily dispersed throughout the genome. This problem is most significant for younger TEs; older elements have accumulated nucleotide substitutions over millions of years that can differentiate them and give rise to uniquely aligning TE reads (Giordano et al. 2007). Because of these barriers, conventional RNA-seq analyses of TEs have either discarded multi-mapping alignments (Chuong et al. 2013) or combined TE expression to the subfamily level (Criscione et al. 2014; Jin et al. 2015; Lerat et al. 2016). Other groups have studied active LINE-1s using tailored pipelines, leveraging internal sequence variation and 3’ transcription extensions into unique sequence (Philippe et al. 2016; Deininger et al. 2017; Scott et al. 2016). However, these targeted approaches are unable to provide a comprehensive picture of TE expression.

To analyze global TE expression in conventional RNA-seq experiments, we have developed the Software for Quantifying Interspersed Repeat Elements (SQuIRE). SQuIRE is the first RNA-seq analysis pipeline available to date that quantifies TE expression at the locus level. In addition to RNA-seq providing expression estimations at the TE locus level, SQuIRE quantifies expression at the subfamily level and performs differential expression analyses on TEs and genes. We benchmark our pipeline using both simulated and experimental datasets and compare its performance against other software pipelines designed to quantify TE expression (Criscione et al. 2014; Jin et al. 2015; Lerat et al. 2016). SQuIRE provides a suite of tools to ensure the pipeline is user-friendly, reproducible, and broadly applicable.

## Results

### SQuIRE Overview

SQuIRE provides a suite of tools for analyzing transposable element (TE) expression in RNA-seq data (Fig. 1). SQuIRE’s tools can be organized into four stages: *1) Preparation, 2) Quantification, 3) Analysis* and *4) Follow-up*. In the *Preparation* stage, **Fetch** downloads requisite annotation files for any species with assembled genomes available on University of California Santa Cruz (UCSC) Genome Browser (Kent et al. 2002). These annotation files include RefSeq (Pruitt et al. 2014) gene information in BED and GTF format, and RepeatMasker (Smit, AFA, Hubley, R & Green) TE information in a custom format. **Fetch** also creates an index for the aligner STAR (Dobin et al. 2013) from chromosome FASTA files. **Clean** reformats TE annotation information from RepeatMasker into a BED file for downstream analyses. The tools in the *Preparation* stage only need to be run once per genome build. Because there are multiple RNA-seq aligners that can produce different results for TE expression estimation, the *Quantification* stage includes the alignment step **Map** to ensure reproducibility. **Map** aligns RNA-seq data using the STAR aligner with parameters tailored to TEs that allow for multi-mapping reads and discordant alignments. It produces a BAM file. **Count** quantifies TE expression using a SQuIRE-specific algorithm that incorporates both unique and multi-mapping reads. It outputs read counts and fragments per kilobase transcript per million reads (fpkm) for each TE locus, and aggregates TE counts and fpkm for TE subfamilies into a separate file. **Count** also quantifies annotated RefSeq gene expression with the transcript assembler StringTie (Pertea et al. 2015) to output annotated gene expression as fpkm in a GTF file, and as counts in a count table file. In the *Analysis* stage, **Call** performs differential expression analysis for TEs and RefSeq genes with the Bioconductor package DESeq2 (Love et al. 2014; Huber et al. 2015). To allow users to visualize alignments to TEs of interest visualized by the Integrative Genomics Viewer (IGV)(Robinson et al. 2011) or UCSC Genome Browser, the *Follow-up* stage tool **Draw** creates bedgraphs for each sample. **Seek** retrieves sequences for genomic coordinates supplied by the user in FASTA format.

**Figure 1.**
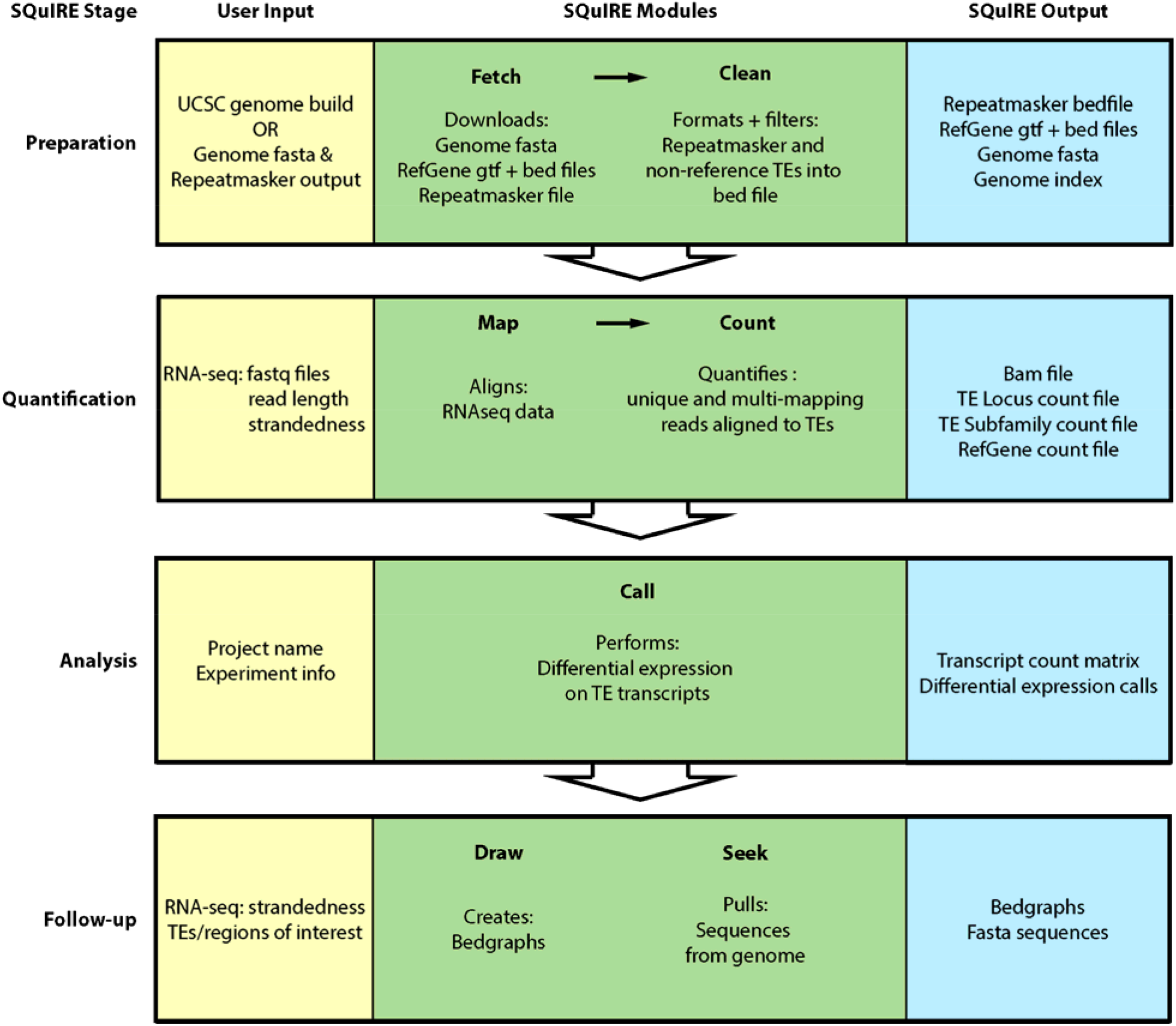
Schematic overview of SQuIRE pipeline.

### Count Algorithm

SQuIRE’s **Count** algorithm addresses a fundamental issue with quantifying reads mapping to TEs: shared sequence identity between TEs from the same subfamily and even superfamily. When a read fragment originating from these non-unique regions is aligned back to the genome, the read may ambiguously map to multiple loci (“multi-mapped reads”). This is not a major problem for older elements that have acquired relatively many nucleotide substitutions, and thus give rise to primarily uniquely aligning reads (“unique reads”). However, TEs from recent genomic insertions that have high sequence similarity to other loci may have few distinguishing nucleotides. Among elements of approximately the same age, relatively shorter TEs also have fewer sequences unique to a locus. Thus, discarding or misattributing multi-mapped reads can result in underestimation of TE expression.

Previous TE RNA-seq analysis pipelines have been able to quantify TE expression at subfamily-level resolution. The software RepEnrich (Criscione et al. 2014) “rescued” multi-mapping reads by re-aligning them to repetitive element pseudogenome assemblies of TE loci and assigning a fractional value inversely proportional to the number of subfamilies to which each read aligned. These multi-mapped fractions were combined with counts of unique reads aligned to each subfamily. This approach was an advance in that it used information from multi-mapped reads. However, this method results in assigning fractions that are proportional to the number of subfamilies that share the multi-mapped read’s sequence, rather than each subfamily’s approximate expression level. TEtranscripts (Jin et al. 2015) expanded on this rescue method by assigning an initial fractional value inversely proportional to the number of TE loci (not subfamilies) to which each read aligned. This initial fractional value was then used in an expectation-maximization (EM) algorithm, which iteratively re-distributes fractions of a multi-mapping read among loci (E-step) in proportion to their relative multi-mapped read abundance estimated from a previous step (M-step). The total of multi-mapped reads and unique reads for each loci are then summed by subfamily. However, in excluding unique reads from the EM algorithm, TEtranscripts does not incorporate empirical high-confidence data to infer TE expression levels from unique TE alignments. Furthermore, in calculating the relative expression level of multi-mapped reads, TEtranscripts normalizes read counts based on annotated coordinates from RepeatMasker. This underestimates TE expression levels for transcripts shorter than the annotated genomic length. TEtranscripts then sums the unique and multi-mapping counts for each subfamily.

In order to accurately quantify TE RNA expression at locus resolution, **Count** builds on these previous methods by leveraging unique read alignments to each TE to assign fractions of multi-mapping reads (Fig. 2). First, **Count** identifies reads that map to TEs (by at least 50% of the read length) and labels them as “unique reads” or “multi-mapped reads.” Second, **Count** assigns fractions of a read to each TE as a function of the probability that the TE gave rise to that read. Uniquely aligning reads are considered certain (e.g., probability = 100%, count = 1). **Count** initially assigns fractions of multi-mapping reads to TEs in proportion to their relative expression as indicated by unique read alignments. In doing so, **Count** also considers that TEs have varying uniquely alignable sequence lengths. To mitigate bias against the *n* number of TEs without uniquely aligning reads, these TEs receive fractions inversely proportional to the number of loci (*N)* to which each read aligned. Then **Count** assigns the remainder 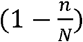 to the TEs with unique reads. To account for TEs that have fewer unique counts due to having less unique sequence, **Count** normalizes each unique count (*C_U_*) to the number of individual unique read start positions, or each TE’s uniquely alignable length (*L_U_*). Among all TEs to which a multi-mapping read aligned, the TEs with unique reads (*s ∊ T*) are compared with each other. A fraction of a read is assigned to each TE in proportion to the contribution of the normalized unique count 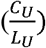 to the combined normalized unique count of all of the TEs being compared 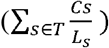. (Equation 1). The sum of unique counts and multi-mapped read fractions for each TE provides an initial estimate of TE read abundance based on empirically obtained unique read counts and uniquely alignable sequence.

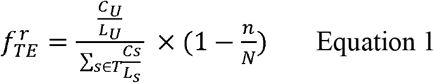

**Figure 2.**
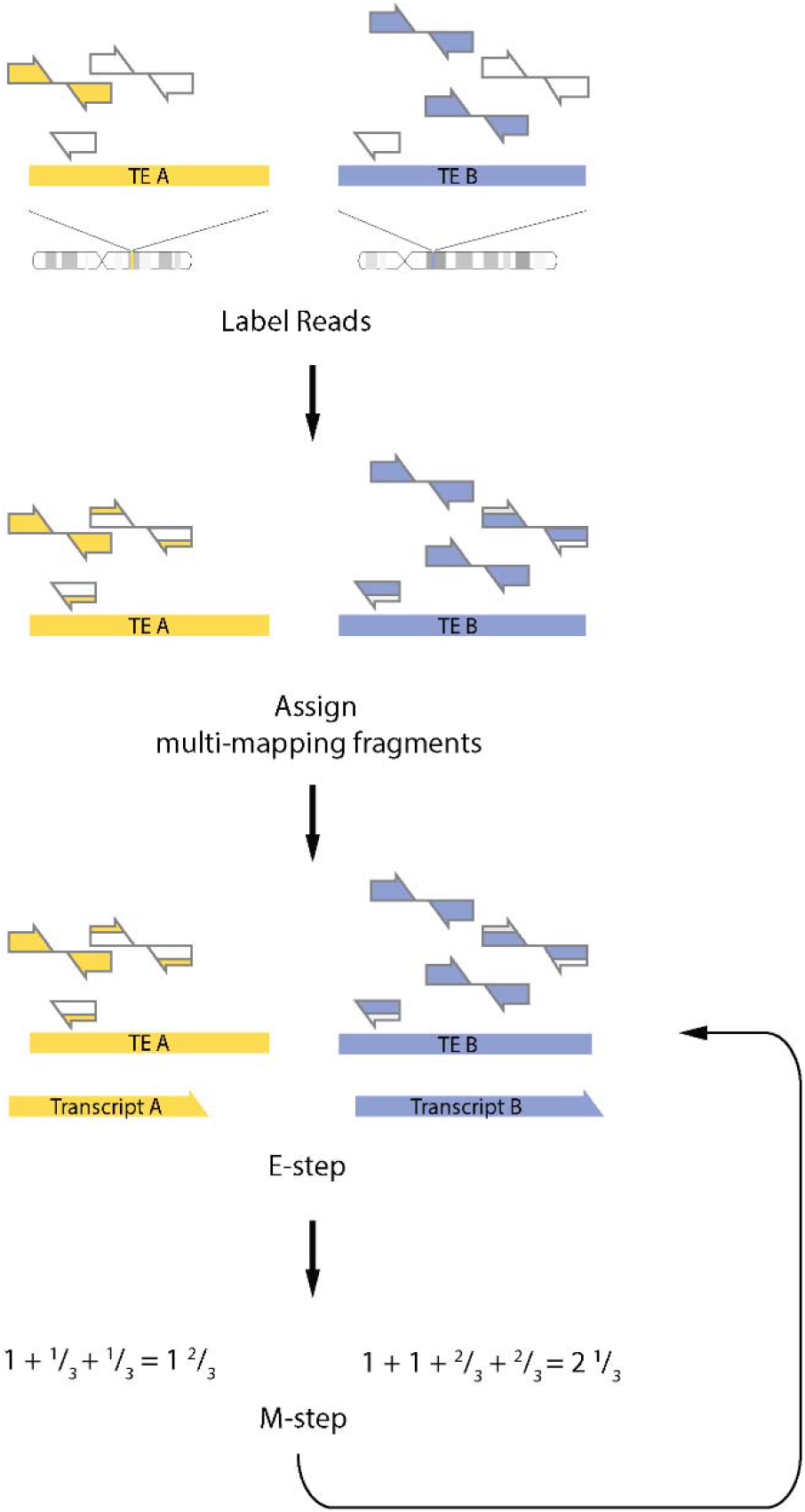
Schematic representation of the SQuIRE **Count** algorithm. First, **Count** labels reads as unique (filled arrows) or multi-mapping (empty arrows). Second, **Count** assigns fractions of multi-mapping reads in proportion to the normalized unique read expression of each TE. The partially filled arrows reflect the proportion of the read assigned to the TE of the corresponding color. Then, **Count** runs an Expectation-Maximization loop that estimates transcript length and reassigns multi-mapping reads for each TE (E-step), then re-estimates total read counts (M-step) until convergence.

Multi-mapping read assignment to TEs without unique reads is thus initially based on the numbers of valid alignments for each read. Count next refines this initial assignment by redistributing multi-mapping read fractions in proportion to estimated TE expression. To estimate expression, **Count** uses the a TE’s total read count (*C_TE_* = unique read counts + multi-mapped fractions from the previous step) normalized by the effective transcript length 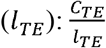. The effective transcript length *l_TE_* is calculated as the estimated transcript length *L_TE_* subtracted by the average fragment length aligned to that TE + 1 (*l_TE_* – *l_TE_* - *l_avg_* + 1), as described previously (Li et al. 2010). All of the TEs to which a multi-mapping read aligned (s *∊ T*) are compared with each other. A fraction of a read is assigned to each TE in proportion to the relative normalized total count 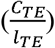 compared to the combined normalized total count of all of the TEs being compared 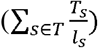, as shown in Equation 2. **Count** assumes this value is proportional to the probability that the TE gave rise to the multi-mapping read, and assigns that fraction of a read count to the TE. Because TEs with a count fraction of less than 1 have a low probability of giving rise to any read, those TEs are assigned a count fraction of 0.

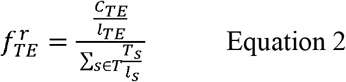

After the total counts (unique and multi-mapped) of each TE are re-calculated, multi-mapped reads can be re-assigned in subsequent iterations of expectation (assigning multi-mapped read fractions to TEs) and maximization (summation of unique and multi-mapped fraction counts). These iterations can be repeated until a given iteration number set by the user or until the TE counts converge (“auto”, when all of the TEs with ≥ 10 counts change by < 1%). An example of **Count** output is provided in Supplemental Table S1. Further details of the **Count** algorithm are in Supplemental Methods.

### Assessing Count Accuracy in simulated data

To test the performance of **Count**, we simulated RNA-seq data from 100,000 randomly selected TEs from the human GRCh38/hg38 (hg38) RepeatMasker annotation (see Methods). TEs were simulated with read coverages of ranging from 2-4000X and simulated counts ranging from 2-4588.We first evaluated accuracy by how closely SQuIRE **Count** output corresponded to the simulated read counts (i.e., % Observed/Expected). However, using this calculation is not meaningful for TEs with low simulated counts: a TE with 0 counts gives an infinite value, and a reported count of 1 for a TE with 2 simulated reads gives a low 50% Observed/Expected. Thus, we were primarily interested in ‘expressed’ simulated TEs, considering only the 99,567 TEs with at least 10 simulated reads. Second, we evaluated SQuIRE by how often it correctly detected simulated TE expression (i.e., true positives) or misreported unexpressed TEs (i.e., false positives).

To test how well SQuIRE performed leveraging only uniquely aligning read information, we first evaluated the % Observed/Expected of TE counts with 0 E-M iterations. We found that SQuIRE accurately assigned read counts to most TEs, with a mean % Observed/Expected of 98.79% (Supplemental Fig. S1). We predicted that this accuracy would be lower for TEs with less uniquely alignable sequence. Indeed, SQuIRE was less accurate for elements with less than 10% divergence (mean of 77.35 % Observed/Expected). The most frequently retrotranspositionally active TEs (i.e., *Alu*Ya5, *Alu*Ya8, *Alu*Yb8, *Alu*Yb9, and L1HS) had counts ranging from 48-70% Observed/Expected, with a range of 79-92% Observed/Expected at the subfamily level (Supplemental Table S2). This illustrates that even without the EM-algorithm, SQuIRE is sensitive for highly homologous subfamilies at the subfamily level.

Given the low recovery of simulated counts for younger elements when relying solely on uniquely aligning reads, we next evaluated how much adding the EM-algorithm improved **Count’s** performance. We anticipated that the counts for most TEs would not change, but that younger elements with less divergence would have improved recovery of simulated reads. Indeed, the overall % Observed/Expected counts of TE loci increased only slightly by 0.14% to a total of 98.93%. However, the change in % Observed/Expected of TEs was much greater for the most homologous active elements, improving by 20.47% for young *Alu* elements and by 21.1% for L1HS loci (Fig. 3). At the subfamily level, the % Observed/Expected of active TEs was improved by 8.1% for young *Alu* elements and by 2.2% for L1HS (Supplemental Table S2). Using updated transcript information in the EM-algorithm is thus particularly useful for TE biologists interested in younger elements that have previously been problematic to quantify by RNA-seq.

**Figure 3.**
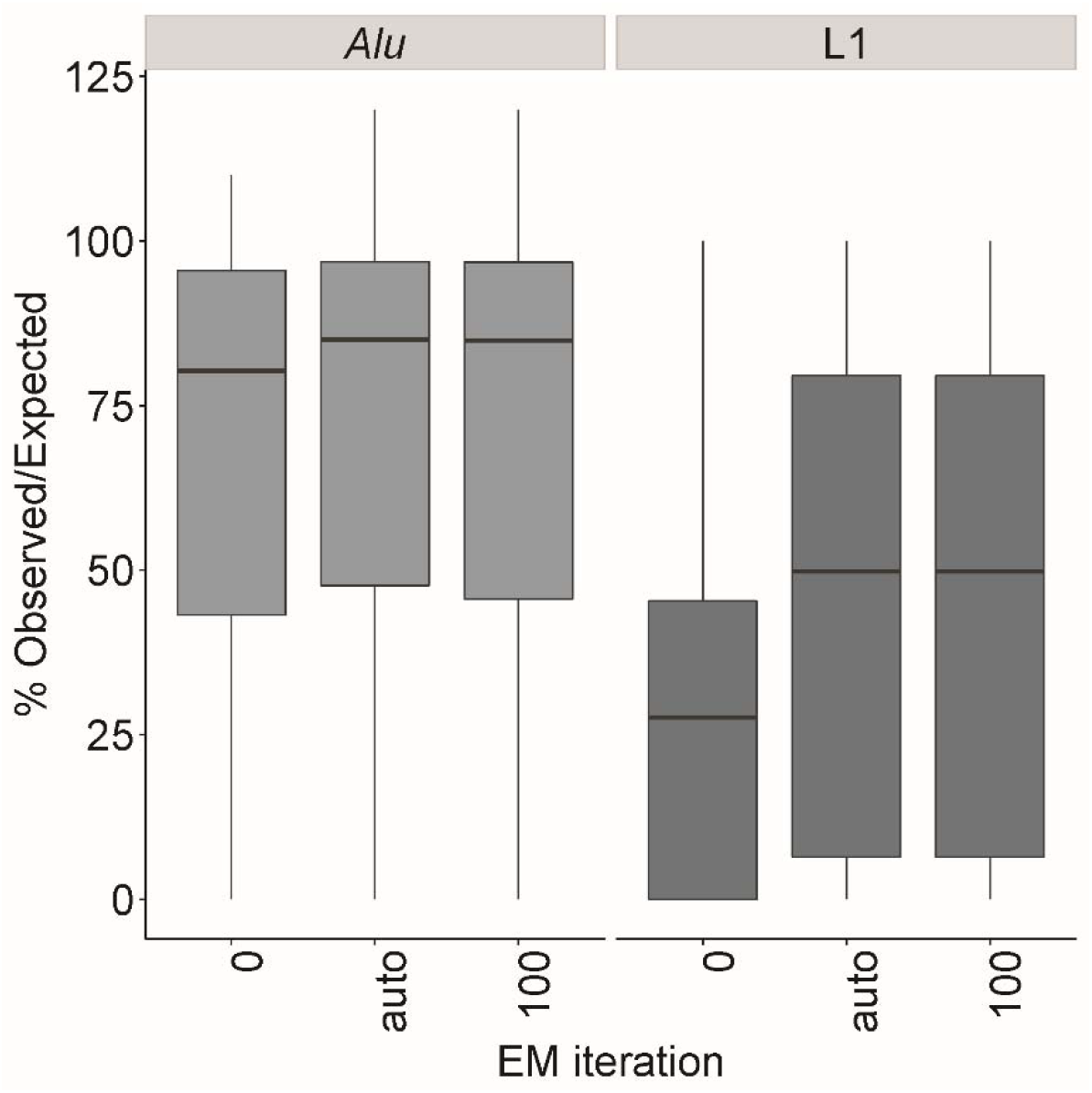
Running EM iterations improves the % Observed/Expected for SQuIRE **Count** for the frequently retrotranspositionally active *Alu* (*Alu*Ya5, *Alu*Ya8, *Alu*Yb8, *Alu*Yb9) and L1 (L1HS) subfamilies compared to no EM iterations (i=0), and does not degrade with increasing iterations (i=100). By default (i=”auto”), SQuIRE **Count** continues the EM-algorithm until each TE with more than 10 reported read counts changes by less than 1%.

We also wanted to evaluate SQuIRE’s ability to distinguish whether a TE is expressed or not. To examine how well **Count** detected expressed TEs, we calculated the true positive rate (TPR) as the percentage of TEs with at least 10 simulated reads that SQuIRE also reported to have ≥ 10 counts. Conversely, we evaluated how often SQuIRE falsely reports TE expression by calculating the positive predictive value (PPV) as the percentage of TEs with ≥ 10 reported counts that were in fact simulated to have ≥10 reads. The true negative rate, or how often SQuIRE correctly reports that a TE is *not* expressed, is less informative for evaluating TE estimation accuracy because the number of TEs in the hg38 genome is so high (>4 million TEs) that the true negative value would outweigh the false positive value (Saito and Rehmsmeier 2015). Overall, SQuIRE had both a high TPR of 98.5% and high PPV of 99.4%. These values were lower for frequently retrotranspositionally active *Alu*s (TPR=68.75-83.33%, PPV= 64.29- 100% ) and L1HS (TPR=100%, PPV=62.86%) using only unique reads for TE expression estimation (Supplemental Table S3). However, using the EM algorithm improved the TPR for *Alus* (TPR=85.22%- 100%) by reducing false negative reports and the PPV for L1HS (PPV=78.57%) by reducing false positives.

### Endogenous LINE-1 detection with Count

To assess **Count’s** ability to detect endogenous LINE-1 expression using biological data, we evaluated the expression level of L1 at loci previously characterized by other methods. Because L1s often become 5’ truncated upon insertion (Perepelitsa-Belancio and Deininger 2003), Deininger et al. performed 5’ rapid amplification of cDNA ends (RACE) on cytoplasmic HEK293 RNA to enrich for full-length L1 RNA. They also performed RNA-seq on polyA-selected cytoplasmic HEK293 RNA to identify L1 loci that have downstream polyadenylation signal. We filtered their findings for L1 loci that had > 5 mapped RNA-seq reads from both 5’RACE and poly-A selected RNA libraries (Deininger et al. 2017) to compare with SQuIRE. We then examined the expression reported by SQuIRE at these 33 loci in paired-end, total RNA from HEK293T cells (GSE113960). We found that 31 (93.4%) had > 10 SQuIRE read counts, confirming their expression (Supplemental Table S4). This suggests that **Count** can detect L1 expression in RNA-seq libraries that are not enriched for L1 loci.

Only a subset of the L1s evaluated by Deininger et al. belonged to L1HS, the youngest family of L1s. Because L1HS loci can be retrotranspositionally active, they can generate insertions that are polymorphic or novel compared to the the reference human RepeatMasker annotation. Reads from TE insertions that are not present in the RepeatMasker annotation can be misattributed to unexpressed, fixed TEs, which can result in “false positive” reports of expression at silent loci. To test how this affects **Count,** we transfected HEK293T cells with an empty pCEP4 plasmid or with a plasmid containing L1RP, an L1HS with known retrotransposition activity (Schwahn et al. 1998; Kimberland et al. 1999). The transfection of L1RP resulted in increased L1HS-aligning reads (254,681 reads) compared to L1HS loci in L1RP-negative cells (2,671 reads) (Supplemental Fig. S2). The differences in L1HS expression in L1RP-transfected cells was higher than what we would expect from endogenous, polymorphic insertions based on previous estimates of polymorphic and fixed L1HS expression in HEK293T cells using unique reads within 1kb downstream of L1HS loci (Philippe et al. 2016). Because Philippe et al. suggested that polymorphic L1HS insertions were transcribed at levels similar to fixed full-length L1HS loci, we sought to mimic polymorphic L1HS expression levels more consistent with previously reported levels. To determine comparable fixed L1HS expression levels in our control HEK293T RNA-seq data, we examined the **Count** output at loci with reported expression by Phillipe et al. (145 read counts). We then downsampled the L1RP-aligning reads from L1RP transfected HEK293T cells to a similar number (153 reads). To simulate a range of polymorphic L1HS expression levels, we also downsampled RNA-seq reads that aligned to the L1RP plasmid to 2X and 20X the fixed active L1HS expression level (302 and 3,091 reads). For these downsampled reads, we identified their other, off-target alignments to the reference genome. To control for potential biological effects of L1RP transfection on TE counts, we ‘spiked in’ these downsampled reads from L1RP-transfected cells into RNA-seq data from HEK293T cells transfected with an empty pCEP4 plasmid. We then calculated the number of false positive L1 loci that became ‘expressed’ with > 10 counts after the *in silico* spike-in. We focused on the 3 youngest L1 subfamilies that share the greatest homology with the L1RP sequence (i.e., L1HS or L1PA1, L1PA2, and L1PA3) (Smit et al. 1995; Boissinot et al. 2000; Lee et al. 2007) and compared their false positive rates to older L1 loci (Fig. 4). When the alignments of 153 reads were spiked in, we found that the false positive rate (FPR) of the youngest L1 subfamilies were comparable to each other, ranging from 34-38%. However, as the spiked in alignments increased to 302 and 3091 reads, the FPR increased for L1HS to 50.68% but not the other subfamilies. This indicates that polymorphic L1HS expression primarily affects the alignments to L1HS loci, and not the loci of closely related subfamilies.

**Figure 4.**
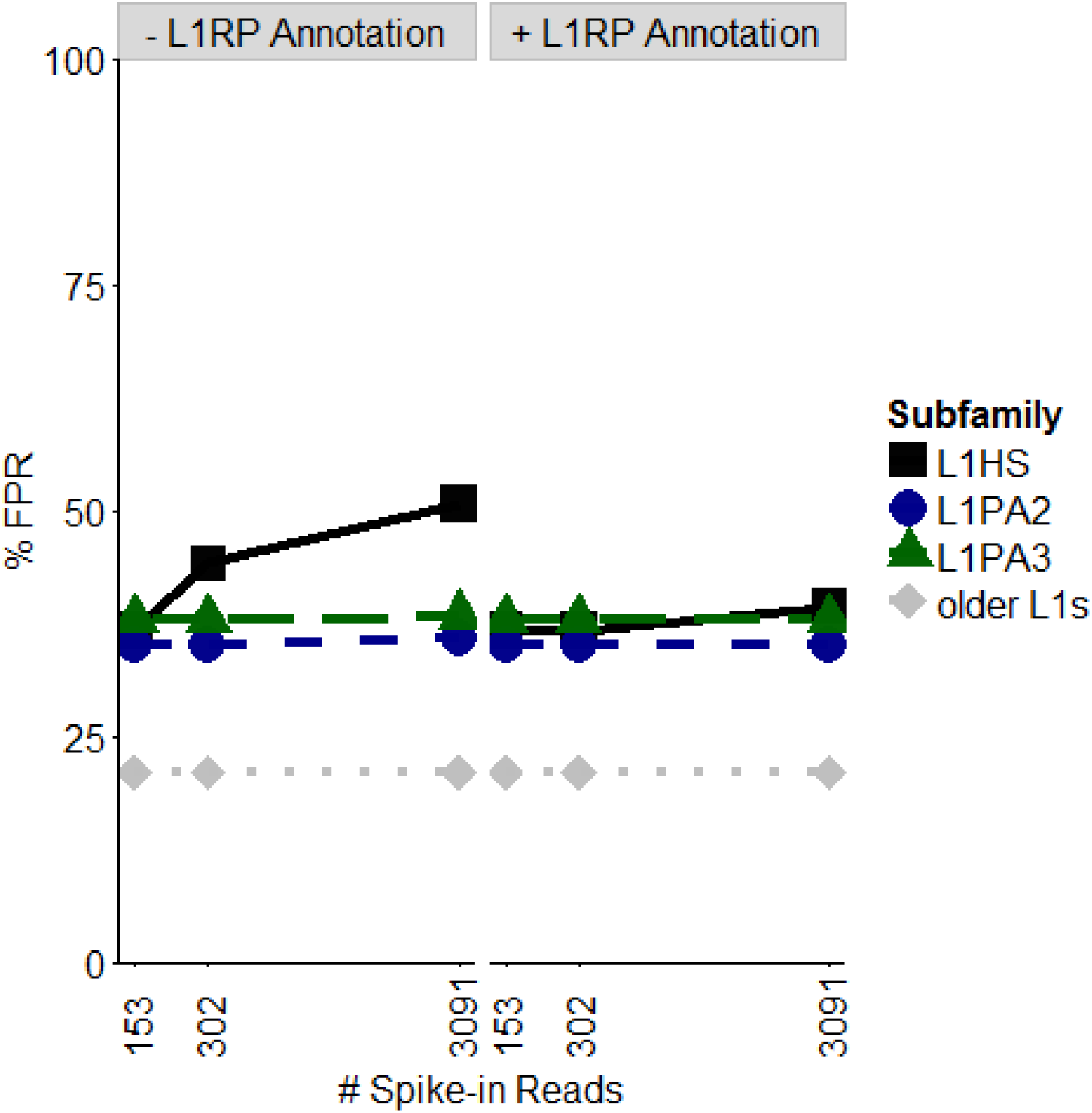
False positive rate (FPR) of L1 loci expression in HEK293T cells when spiking in L1RP-aligning reads. False positive expression is implicated a locus that previously had <10 reads has ≥ 10 reads after spike-in. % FPR is the percentage of loci with false positive loci relative to the total number of loci with ≥ 10 SQuIRE read counts. The number of spike-in reads (153, 302, 3091) represents 1X, 2X and 20X predicted endogenous polymorphic L1HS expression levels based on findings from Phillipe et al. 2016. The FPR is robust for older L1 subfamilies with increased spike-in reads. The addition of L1RP annotation in a non-reference table reduces the change in false positive rate for L1HS after increasing spike-in reads.

L1-mapping methods (Upton et al. 2015; Rodić et al. 2015; Iskow et al. 2010; Ewing et al. 2010) and TE insertion detection software for whole genome sequencing (Gardner et al. 2017; Lee et al. 2012; Keane et al. 2013; Stewart et al. 2011; Sudmant et al. 2015; Ewing et al. 2011) can identify locations of non-reference TE insertions. Validating these insertions by PCR and Sanger sequencing can provide not only unique sequence flanking the insertion but potentially also the TE sequence. Users can input a custom table to SQuIRE **Map** and **Clean** (Supplemental Table S5) to add non-reference TEs and their flanking sequence to the alignment index and RepeatMasker BED file. We evaluated how incorporating the non-reference table containing information about the L1RP plasmid affected the FPR in HEK293T cell data. We found that the FPR for L1HS only increased from 36.67% with 153 reads spiked in to 39.34% with 3091 reads spiked in. Thus, adding L1RP information improved **Count’s** accuracy at higher L1RP *in silico* expression levels.

### Comparison to other software

Currently published TE analysis software include RepEnrich, TEtranscripts, and TETools (Criscione et al. 2014; Jin et al. 2015; Lerat et al. 2016). We used the simulated hg38 TE data described above to compare the recovery of simulated reads to the correct subfamily among TE quantification software (% Observed/Expected). For mapping, we ran each software’s recommended aligner: STAR (used by SQuIRE and TEtranscripts), Bowtie 2 (used by TETools), and Bowtie 1 (used by RepEnrich). We found that SQuIRE (99.86% ±1.46 %), TETools (100.14 ± 2.21%), and TEtranscripts (95.89 ± 16.41%) had comparable % Observed/Expected rates (Supplemental Fig. S3). In contrast, RepEnrich (108.77 ± 40.67%) was less accurate in terms of % Observed/Expected. This is likely attributable to RepEnrich’s recommended use of Bowtie 1, which discards discordant reads and limits the number of attempts to align both paired-end mates to repetitive regions. To support this, we compared how often each aligner mapped a uniquely aligning simulated read to the correct location. We indeed found that Bowtie 1 failed to report unique reads more often in a paired-end library compared to single-end (Supplemental Table S6).

To compare SQuIRE to other TE analysis tools with biological data, we ran each pipeline on publically available adult C57Bl/6 mouse tissue RNA-seq data (Brawand et al. 2011) using GRCm38/mm10 (mm10) TE annotation. We compared the expression of subfamilies in testis compared to pooled data from brain, heart, kidney, and liver tissues. To independently evaluate the fold-changes of TE RNA between testis and somatic tissues, we also used our previously published adult C57Bl/6 mouse Nanostring results (Gnanakkan et al. 2013). Unlike RNA-seq analysis, which infers transcript levels by counting reads, Nanostring uses uniquely mapping probes to capture and count RNA molecules. We compared the Nanostring log_2_ fold changes (log_2_FC) of TE subfamily expression in testis and pooled somatic tissue to the log_2_FC values found by SQuIRE, RepEnrich, TEtranscripts, and TETools (Supplemental Fig. S4). We first looked at how often the direction of fold change corresponded between each tool and Nanostring. For the 16 subfamilies queried, SQuIRE and TETools shared the same direction of fold change as Nanostring more often than the other tools (SQuIRE: 12, TETools: 12, TEtranscripts: 9, RepEnrich: 8). Moreover, compared to TETools, SQuIRE reported log2FC values closer to the expected values from Nanostring (mean absolute differences in log2FC from Nanostring– SQuIRE: 0.965, TETools: 1.34, TEtranscripts: 1.16, RepEnrich: 1.11).

With SQuIRE, we can closely examine the mouse RNA-seq data at the locus level. For the 16 subfamilies analyzed by Nanostring and the TE analysis tools, we found that the reported subfamily-level expression could be attributed to fewer than 7% of each subfamily’s loci (Supplemental Fig. S5). This suggests that regulation of TE transcription is not necessarily shared across all TEs from the same subfamily. On the other hand, whereas the other subfamilies studied by Nanostring have only 1-4 significantly differentially expressed loci (log2FC >1, padj < 0.05), the IAPLTR3 subfamily has 11 loci that are all differentially expressed in testis compared to somatic tissues (Fig. 5A). To test whether this was an enrichment relative to the representation of IAPLTR3 in the mouse genome, we performed a Fisher’s exact test and found that IAPLTR3 loci were 10-fold more likely than expected to be differentially expressed in testis (OR: 10.56, 95%CI: 5.25-18.97, padj < 1.61 e-08). This suggests that a subset of TE locus expression may still be impacted by subfamily-specific regulation.

**Figure 5.**
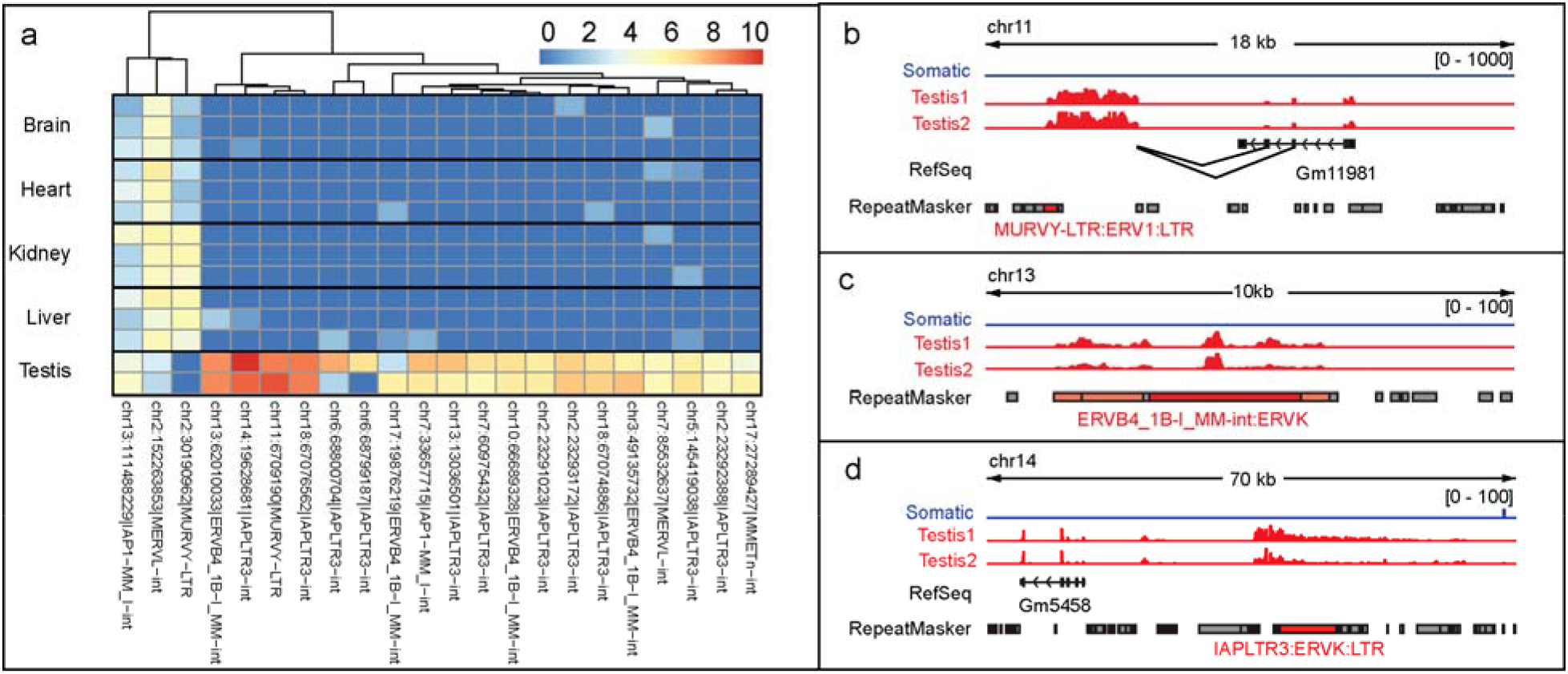
Differentially expressed TE loci belonging to subfamilies previously analyzed by Nanostring a. The X-axis represents replicates of somatic and testis tissue samples from adult C57Bl/6 mouse. The Y-axis represents differentially expressed TE loci. The heatmap colors repres the log2 of total read counts +1 for each TE locus. b-d. Examples of intergenic TE loci differentially expressed in testis compared to somatic tissues. Tracks from brain, heart, kidney and liver replicates were collapsed into a single track. The scales of count expression are shown in brackets. The RefSeq track represents annotated genes. The RepeatMasker track represents transposable elements annotated in the reference genome. Transposable elements colored in red belong to the subfamily indicated; dark red indicates that that RepeatMasker entry meets significant differential expression thresholds (log2FC > 2, padj < 0.05).

To further investigate the interplay between genomic context and TE subfamily, we identified the closest genes to each differentially expressed locus and clustered the loci by their expression levels, as shown in Figure 5A. We found a cluster of 3 loci exhibiting broad expression across somatic tissues from the IAP1, MERVL, and MURVY LTR retrotransposon subfamilies. When we examined the genomic context of these 3 loci, we found that all were located within genes with known broad tissue expression (*Gpbp1*, *Csnk2a1*, *Kyat1*, respectively) (Yue et al. 2014), with examples shown in Supplemental Figure S6. Another locus from the MURVY subfamily is in a cluster of TEs exhibiting high testis-restricted expression. In examining the transcript overlapping the MURVY locus, we see that the transcript initiates outside of the locus and find that the transcript is an alternative splicing isoform with splice donors from the third and fourth exons of a gene ~5kb away (Fig. 5B). The gene, *Gm11981*, is a long noncoding RNA (lncRNA) known to exhibit testis-restricted expression (Yue et al. 2014). The different MURVY-containing transcripts illustrate how the relationship between TE expression and neighboring transcription can vary across loci from the same subfamily. We also examined ERVB4-1B and IAPLTR3, the two LTR retrotransposon subfamilies that exhibited the highest fold change by Nanostring. These subfamilies were represented in the high-expressing, tissue-restricted loci cluster (Fig. 5A). While the transcription of the ERVB4-1B locus on chr13 did not extend beyond annotations for that subfamily (Fig. 5C), the IAPLTR3 loci on chr14 (Fig. 5D) and chr18 are part of longer transcripts that initiate outside of the annotated TE. Unlike the MURVY locus on chr11, there is no evidence of splicing into the IAPLTR and ERVB4-1B loci. Thus, TEs from different subfamilies may be subject to different mechanisms of transcriptional regulation as evidenced by expression within different transcript structures. Altogether, this stresses the utility of using SQuIRE to analyze TE transcription at the locus level.

### Benchmarking for SQuIRE’s Memory Usage and Running Time

To benchmark SQuIRE’s memory usage and running time for RNA-seq data of different sequencing depths, we subset the high-depth (mean 263 million reads across 8 lanes) HEK293T cell line RNA-seq data into 1, 2, and 3-lane libraries with a mean sequencing depth of 32, 65, and 98 million reads. We evaluated the speed and memory performance of each *Quantification* and *Analysis* stage tool for each sequencing depth (Fig. 6) using 8 parallel threads and 64 Gb of available memory. We found that sequencing depth had the greatest effect on **Count**, taking 8.6 hours to complete the 3-lane library compared to 2.4 hours for the 1 lane library. The other tools took much less time and were less affected by sequencing depth. **Map** took 1-2 hours for the different libraries. **Call** running time was also independent of library size, but it was greater when including all TE counts (10 minutes) compared to subfamily counts (2 minutes). We found that the memory usage of each tool was largely independent of sequencing depth, taking between 39-40Gb of Memory for **Map**, 30-32Gb for **Count,** and 7-8Gb for **Call**.

**Figure 6.**
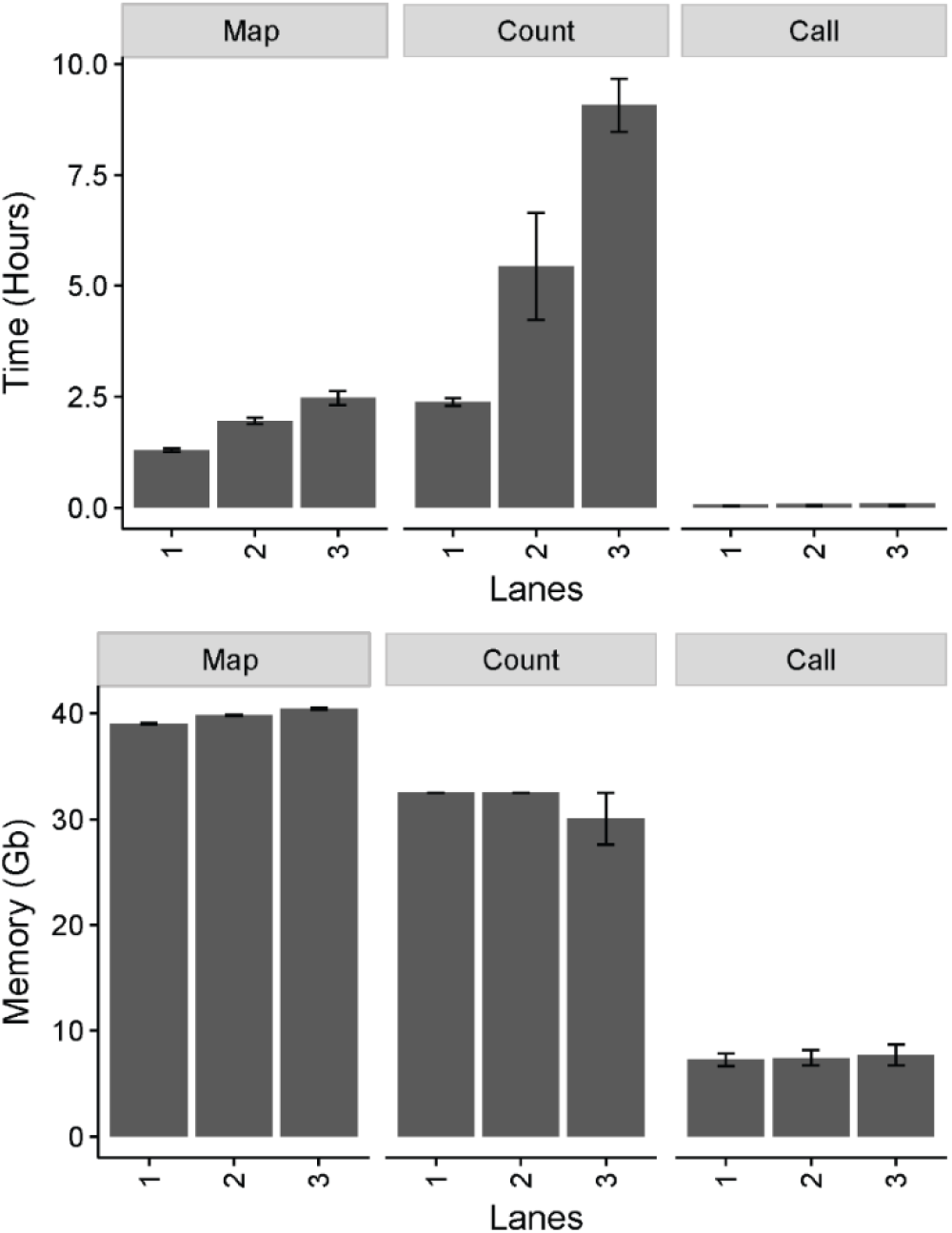
Usage data for the main modules of SQuIRE. Time (Hours) and Memory for SQuIRE **Count, Map** and **Call**. Mean library sizes for RNA seq data were 1 lane= 32,912,528 reads, 2 lanes= 65,573,850 reads, 3 lanes= 98,757,439 reads.

### Implementation

Our efforts at making SQuIRE easy to use has resulted in a simple installation process in which the user can copy and paste lines of code to install all prerequisite software and set up SQuIRE (Table 1). In addition, SQuIRE is the only program that downloads reference annotation for assembled genomes available on UCSC, allowing it to be easily adaptable to a variety of species. For genomes from non-model organisms or organism strains with high divergence from the reference annotation, SQuIRE can also use RepeatMasker software output for even wider compatibility. To ensure that the pipeline is streamlined and that the outputs are reproducible, SQuIRE also implements alignment and differential expression for the user. In making SQuIRE as user-friendly as possible, we intend to improve the reproducibility of bioinformatics in the TE field.

**Table 1.** Feature comparison of RNA-seq Analysis tools for TEs.

|  | SQuIRE | RepEnrich | TETranscripts | TETools |
| --- | --- | --- | --- | --- |
| Provides Locus-level TE RNA quantification | **YES** | -- | -- | -- |
| Provides TE transcript information | **YES** | -- | -- | -- |
| Copy-and-paste installation | **YES** | -- | -- | -- |
| Provides prerequisite annotation files for any species | **YES** | -- | -- | -- |
| Can incorporate non-reference TEs | **YES** | -- | -- | **YES** |
| Performs alignment | **YES** – uses STAR | Recommends Bowtie 1 | Recommends STAR | **YES** – uses Bowtie 1 or Bowtie 2 |
| Uses genome for alignment | **YES** | **YES** - Genome + TE pseudogenome | **YES** | -- |
| Provides gene expression quantification | **YES** | -- | **YES** | -- |
| Performs differential expression | **YES** | -- | **YES** | **YES** |

## Discussion

We have developed Software for Quantifying Interspersed Repeat Expression (SQuIRE) to characterize TE expression using RNA-seq data. TEs are highly repeated in the genome, which can pose challenges for mapping reads unambiguously to specific transcribed loci. SQuIRE is the first RNA-seq analysis software that provides locus-specific TE expression quantification while also outputting subfamily-level expression estimates (Table 1). Our approach uses unambiguously mapping reads and an Expectation-Maximization algorithm to estimate levels of TE transcripts. SQuIRE additionally provides information on the structure of each TE transcript, which can be shorter or longer, sense or antisense compared to the annotated repeat. We have shown that SQuIRE can correctly attribute a high percentage of reads originating from TEs using simulated data. Although this percentage is lower for frequently retrotranspositionally active, less divergent TEs (e.g., *Alu*Ya5, *Alu*Ya8, *Alu*Yb8, *Alu*Yb9, L1HS), we found that implementation of an Expectation-Maximization (EM) algorithm (Jin et al. 2015; Li and Dewey 2011) improves accuracy and lowers both false positive and false negative estimations of whether a TE is expressed. This finding also holds in biological settings, where SQuIRE is able to correctly identify instances of full-length L1 expression in total RNA RNA-seq data from cell lines wherein previous studies had identified these loci using a combination of 5’RACE and 3’ primer extension methods (Deininger et al. 2017). This confirms that SQuIRE can detect the expression of TEs in the reference genome that have in the past been problematic for global TE RNA expression analysis.

The ongoing activity of TEs also results in a significant number of mobile element insertion variants (MEI) (Beck et al. 2010; Sudmant et al. 2015; Stewart et al. 2011). Numerous commonly occurring structural variants owed to retrotransposition are missing in reference genome assemblies. SQuIRE provides users with two options to query transcription of these repeats. First, it can detect their transcription at the subfamily level. We have shown that SQuIRE can detect expression of L1HS elements when we express an ectopic sequence. It maintains a low false positive rate of misattributing these reads to endogenous L1HS loci. Thus, SQuIRE can be useful for detecting altered regulation of young TE subfamilies even when specific loci that are expressed are unknown. Secondly, SQuIRE can use sequences of known, non-reference TE insertion polymorphisms to detect locus-specific expression when these are available. For example, in the human genome, L1HS element sites and sequences can be obtained by targeted TE insertion mapping (Upton et al. 2015; Rodić et al. 2015; Iskow et al. 2010; Ewing et al. 2010) or whole genome sequencing (Gardner et al. 2017; Lee et al. 2012; Keane et al. 2013; Ewing et al. 2011). Polymorphic TE insertions have been reported to databases such as euL1db (Mir et al. 2015), dbRIP (Wang et al. 2006) and 1000 Genomes Project (Sudmant et al. 2015). If the polymorphic insertions have been verified and sequenced in the user’s samples, SQuIRE is capable of incorporating user-provided, non-reference TE sequence to estimate TE expression at these loci. This may be a useful feature for understanding functional consequences of these insertion variants (Payer et al. 2017).

The SQuIRE algorithm builds on strategies used by previous TE analysis software (Criscione et al. 2014; Jin et al. 2015; Lerat et al. 2016). Here, we show that SQuIRE provides additional features and improves on the accuracy of these methods, as assessed using both simulated reads and orthogonal approaches to measure log_2_ fold changes in mouse tissue comparisons. Our findings suggest that important biologic insights can be gained by examining TE transcription at the locus level.

To date, locus-specific studies of TE expression and activity have mostly focused on identifying transcriptionally and retrotranspositionally active L1s in the human genome (Deininger et al. 2017; Philippe et al. 2016; Scott et al. 2016; Brouha et al. 2003; Beck et al. 2010; Tubio et al. 2014; Pitkänen et al. 2014). In applying SQuIRE to study locus-specific TE expression genome-wide in mouse tissues, we can see that this paradigm is not unique to L1s or humans. It seems a very limited subset of TE loci are transcribed with complex patterns of tissue-specific expression. Furthermore, we found that the tissue expression patterns of TE loci were driven by a variety of transcriptome contexts: broadly expressed mRNA transcripts, testis-specific lncRNA and authentic TE ‘unit’ transcripts. How these TEs affect genome regulation remains an open question. Prior to SQuIRE, the inability to map TE expression limited genome-wide analysis of TEs to the effects of *cis*-acting elements on transcriptional (Faulkner et al. 2009; Kalitsis and Saffery 2009; Le et al. 2015; Xie et al. 2013) and post-transcriptional (Stower 2013; Sorek et al. 2002; Ecco et al. 2016; Athanasiadis et al. 2004) regulation. Further, the effects of neighboring genes on TE transcription are not well-understood. In providing locus-level TE transcript estimations, SQuIRE can enable studies that dissect the regulatory impacts of TE and gene expression.

## Methods

### Implementation of STAR aligner in Map

**Map** uses parameters tailored to the alignment of TEs. By default STAR only reports reads that map concordantly and to 10 or fewer locations. **Map** retains more reads mapped to TEs by reporting reads that map to 100 or fewer locations (--outFilterMultimapNmax 100 –winAnchorMultimapNmax 100). For paired-end reads, **Map** also reports paired reads that map discordantly (--chimSegmentMin <read_length>) and single reads with unmapped mates (--outFilterScoreMinOverLread 0.4 – outFilterMatchNminOverLread 0.4). **Map** can incorporate the non-reference TE sequences and generate a FASTA file that STAR adds to the genome index with the option “—genomeFastaFiles <fasta> ”. To provide splicing information to the tools in the *Analysis Stage*, **Map** also uses the UCSC RefSeq gene annotation and assesses reads overlapping splice junctions with the options “—sjdbGTFfile <gtf> -- sjdbOverhang <read_length -1> --twopassMode Basic”. **Map** produces a sorted BAM file that includes intron and splicing information for downstream transcriptome assembly analysis.

### Implementation of StringTie in Count

**Count** runs StringTie (Pertea et al. 2015)using these default settings guided by RefSeq gtf obtained from UCSC with **Fetch. Count** uses the “-e” StringTie option to quantify expression only to annotated transcripts without assembly of novel transcripts. We convert the fpkm values to counts by multiplying the per-exon coverage by exon length normalized by read length.

### DESeq2 Implementation in Call

**Call** incorporates the Bioconductor package DESeq2 (Love et al. 2014; Huber et al. 2015) with its suggested parameters. Users input the sample names and experimental design (ie which samples are treatment or control), which **Call** uses to find **Count** data and create a count matrix for annotated RefSeq genes, StringTie transcripts and TEs. **Call** outputs differential expression tables and generates MA-plots, data quality assessment plots, and volcano plots.

### STAR implementation in Draw

To visualize the distribution of reads across the TE, **Draw** runs STAR (Dobin et al. 2013)with the parameters “–runMode input AlignmentsFromBAM –outWigType bedGraph” to provide visualization of read alignments. It will output bedgraphs of all reads (“multi”) and only uniquely (“unique”) aligning reads. **Draw** also compresses the bedgraphs into bigwig format for IGV (Robinson et al. 2011) and UCSC Genome Browser (Rosenbloom et al. 2014) viewing. If the RNA-seq data is stranded it will output unique and multi bedgraphs for each strand.

### RNA-seq simulation

We randomly selected 100,000 TEs from the hg38 Repeatmasker annotation downloaded by **Fetch**. We limited our list of potential TEs to those included in TEtranscripts (Jin et al. 2015) and RepEnrich (Criscione et al. 2014) to enable comparisons between these different programs. Using the selected TE coordinates we generated a BED file using **Clean** and obtained Fasta sequences using **Seek.** From these TE sequences, we used the Polyester package from Bioconductor (R version 3.4.1,Huber et al. 2015) (Huber et al. 2015)to simulate 100bp, paired-end, stranded RNA-seq reads with normally distributed fragment lengths around a mean of 250bp. We simulated a uniformly distributed sequencing error rate of 0.5%. TEs were simulated with a mean read coverage of 20X, with 250 TEs deviating from that mean between 2-100 fold.

### HEK293T Cell Culture, Transfection and Sequencing

Tet-On HEK293TLD (293T) cells (Taylor et al. 2013) were grown at 37C, 5% CO2 in DMEM with 10% Tet-Free FBS (Takara, Mountain View, CA) and passaged every 3-5 days as needed.

LINE expression constructs were cloned into the pCEP4 backbone (Thermo Fisher Scientific, Waltham, MA) modified to confer puromycin resistance. Plasmids encoded either L1RP (MT302) or had no insert (Taylor et al. 2013). For transfection, 300,000 293T cells were plated in 2 mL volume. 24 hours later, cells were transfected using a cocktail of 2 ug plasmid DNA and 6 uL Fugene HD (Promega), and puromycin was added 24 hours later for a total of 3 days of selection. 500,000 cells were then plated in 3 wells each, and doxycycline was added 2 hours later (final concentration of 1 ug/ml) to induce L1 expression. RNA was collected after 72 hours of L1 expression using the Zymo Quick-RNA MiniPrep kit (Zymo Research, Tustin, CA). The RNA libraries of transfected 293T cells were prepared using the Illumina TruSeq Stranded Total Library Prep Kit with Ribo-Zero Gold (San Diego, CA) to provide stranded, ribosomal RNA depleted RNA. The libraries were sequenced on an Illumina HiSeq 2500, using 6 samples per lane across 8 lanes with paired-end 100bp reads. We generated a mean of 263,127,067 paired reads per sample. The raw sequencing data were deposited to the NCBI Genome Expression Omnibus (GEO) with accession number GSE113960.

### HEK293T Cell RNA-seq Analysis and *In Silico* Spike-in Experiment

For detection of fixed L1 expression identified by Deininger et al. by 5’RACE and poly-A selected RNA sequencing in HEK293 cells, we ran SQuIRE **Map, Count**, and **Call** on HEK293T cell samples transfected with empty L1RP vector (DA5 and DA6). To determine the effect of L1RP transfection on the false positive rate of L1 RNA estimation, we ran **Map** and **Count** on HEK293T cells transfected with L1RP and vector. To simulate the effect of polymorphic TE expression on typical RNA-seq samples, we downsampled a transfected (DA1) and control (DA5) sample to a single lane per sample (average 32 million reads). To identify L1RP aligning reads in the L1RP-transfected cell, we used SAMtools (Li et al. 2009) to identify reads that align to the chromosome construct provided by the non-reference table (Supplemental Table S5). To downsample the L1RP-aligning reads, we used the SAMtools “-s <*INT*.*FRAC*> ” option with 0.01, 1.001, and 3.0004 as inputs. The integer before the decimal indicates the seed value and the number after the decimal indicates the fraction of total alignments desired for subsampling. We then identified all alignments to the genome sharing the same Read IDs as the down-sampled L1RP-aligning reads. We used SAMtools merge to combine the alignments of L1RP-aligning reads with the BAM file of the HEK293T cell sample transfected with empty vector (DA5).

### TE RNA-seq tool Comparison

Adult C57BL/6 mouse RNA-seq data were obtained from GEO with accession number GSE30352. All pipelines were run on a server with a maximum of 128 GB memory available and 8 threads (-p setting).

RepEnrich (Criscione et al. 2014)– We obtained the hg38 annotation for RepeatMasker from the RepEnrich GitHub website. For the mm10 annotation, we obtained the mm10.fa.out.gz RepeatMasker (Smit, AFA, Hubley, R & Green) annotation from the RepeatMasker website. We ran the setup for RepEnrich following instructions from the website for each genome build. We then mapped the data to the genome using Bowtie 1 (Langmead et al. 2009) according to RepEnrich’s instructions to generate separate uniquely mapping sam and multi-mapping read .fastq files. These were then used for the RepEnrich software with the “–pairedend TRUE” parameter for simulated human data, and “—pairedend FALSE” for mouse data.

TETools (Lerat et al. 2016)– We generated rosette files for hg38 and mm10 for TETools by taking the Repeatmasker annotation from **Clean** for the first column and the repeat taxonomy for the second column (subfamily:family:superfamily). We used the BED file from **Clean** with **Seek** to obtain TE FASTA sequences for generation of a pseudogenome for TETools. TETools was run with the “-bowtie2”, “–RNApair” and “–insert 250” parameters for simulated human data and “-bowtie2”,”-insert 76” for mouse data.

TEtranscripts (Jin et al. 2015) –We obtained hg38 and mm10 GTF annotation from the TEtranscripts website. We aligned the data to the genome with STAR using “--winAnchorMultimapNmax 100”,”-- outFilterMultimapNmax 100” parameters for multi-mapping. We then ran TEtranscripts with the “--mode multi” setting to utilize its expectation-maximization algorithm for assigning multi-reads for the resulting SAM file. Since TEtranscripts analyzes TE and gene expression together, we used refGene annotation obtained by SQuIRE **Fetch** for the required gtf file. We used the parameters “--format SAM”, “--mode multi”, “--stranded yes” for simulated human data, and “--format SAM”, “--mode multi”, “--stranded no” for mouse data.

### Aligner Comparison

We ran the aligners Bowtie1 (Langmead et al. 2009), Bowtie2 (Langmead and Salzberg 2012), and STAR (Dobin et al. 2013) on the simulated TE RNA-seq data described above. We set each aligner to output a maximum of 2 valid alignments to quickly identify uniquely aligning reads with the parameter “- m2” for Bowtie 1, “-k2” for Bowtie 2, and “--outSAMmultNmax 2” for STAR. We also ran STAR with the parameters “--outFilterScoreMinOverLread 0.4 --outFilterMatchNminOverLread 0.4 -- chimSegmentMin 100” to allow for discordant alignments, which STAR excludes by default. Bowtie2 reports discordant alignments by default, while Bowtie 1 can only report paired alignments. We used BEDTools (Quinlan and Hall 2010) to intersect the BAM outputs to RepeatMasker annotation to identify the TEs to which the aligners mapped the reads. Reads that only appeared once as “uniquely aligning”.

We assessed whether the mapped TE matched the templating TE for the simulated read to determine if the uniquely aligning reads mapped to the correct location.

### Data Access

The raw sequencing data and SQuIRE Count output for HEK293T cell transfection were deposited to the NCBI Genome Expression Omnibus with accession number GSE113960. SQuIRE was written in Python2 and is available at the website https://github.com/wyang17/SQuIRE and PyPI. It was developed for UNIX environments. We provide step-by-step instructions on our README to install the correct versions of all software. These instructions include using the package manager Conda (conda.io) to download the correct versions of prerequisite software for SQuIRE (e.g., Python, R (R Development Core Team 2011), STAR, BEDTools, StringTie, SAMtools (Li et al. 2009), UCSC tools and Bioconductor packages. The README also instructs users how to create a non-reference table with the exogenous or polymorphic TE sequences and coordinates that they would like to add to the reference genome. Bash scripts to run each tool in the SQuIRE pipeline are also included. Users can fill in crucial experiment information (raw data, read length, paired, strandedness, genome build, sample name and experimental design) into the “arguments.sh” file, which the other scripts reference to run each step with the correct parameters.

## Acknowledgements

We would like to thank Veena Gnanakkan for preparation of C57BL/6 mouse tissue RNA and analysis of Nanostring data. We would like to thank Jane Welch, Paul Schaughency, Shubha Tirumale and Ping Ye for testing SQuIRE. We would like to thank Sibyl Medabalimi for assistance in developing the name of SQuIRE. We would also like to acknowledge the assistance of the NYU Genome Technology Center and Jared Steranka in preparing the RNA for RNA sequencing. This research was supported by National Institutes of Health (NIH) grants R01GM124531 and P50GM107632, and Department of Defense Congressionally Directed Medical Research Program (CDMRP) grant OC120390 (to K.H.B.). W.R.Y. received a Teal Predoctoral Scholar award in association with OC120390.

## Author Contributions

W.R.Y. contributed to study design, programming of SQuIRE pipeline, statistical analysis, and primary authorship and manuscript; D.A. contributed culture and transfection of HEK293T cells, and suggestions for analysis and manuscript; C.N.P. contributed to debugging SQuIRE and development of README for SQuIRE website; L.M.P. & K.H.B. jointly contributed to overall study design, data interpretation and manuscript.

